# Near real-time data on the human neutralizing antibody landscape to influenza virus in summer of 2026 shows antigenic advance of H3N2 subclade K region D mutants and H1N1 D.3.1.1 Sa mutants

**DOI:** 10.64898/2026.09.15.751855

**Authors:** Caroline Kikawa, Andrew Butler, John Huddleston, Sam A. Turner, Heidi Peck, Janet A. Englund, Kirsten Lacombe, Michael Busch, Marion C. Lanteri, Mars Stone, Bryan Spencer, Alexander L. Greninger, Derek J. Smith, Stephanie Wallace, Helen S. Marshall, Shidan Tosif, Scott E. Hensley, Ian G. Barr, Jesse D. Bloom

## Abstract

Human seasonal influenza evolves rapidly, necessitating twice yearly decisions about whether to update the strains in the vaccine. To help inform this decision, we have been using high-throughput sequencing-based neutralization assays to make twice yearly measurements of how recent human sera neutralize current human H3N2 and H1N1 strains. Here we provide the third installment in this series of measurements by reporting 52,268 titers representing neutralization of 148 viral strains by 355 human sera collected between April and August of 2026. Our measurements show that new H3N2 subclade K strains with mutations in antigenic region D and new H1N1 subclade D.3.1.1 strains with mutations in antigenic region Sa (such as G155E) have reduced neutralization by human sera, with notable heterogeneity in the impact of some of these mutations across sera from different individuals. This paper is accompanied by an interactive summary (https://jbloomlab.github.io/flu-seqneut-2026/summary.html) that enables detailed exploration of the results, and all titer data are publicly available for further analysis to aid vaccine antigen selection and studies of viral evolution.

## Introduction

The hemagglutinin (HA) protein of human seasonal influenza evolves rapidly to erode neutralizing antibodies (Smith et al. 2004; Bedford et al. 2014). To keep pace with this evolution, the influenza vaccine is updated regularly, with a decision made twice per year about whether to change the strains in the vaccine to better match circulating strains. Because it takes time to produce and administer the vaccines, the strain update decision is made roughly seven to ten months before the influenza season during which the vaccine will be deployed: vaccine strains are chosen in September for the following year’s Southern Hemisphere influenza season, and in February for the Northern Hemisphere influenza season that begins at the end of the same year (World Health Organization 2026; World Health Organization 2025). This gap between the timing of the vaccine update decision and the influenza season requires forecasting which strains will dominate nearly a year in the future.

The influenza strains that spread in the human population tend to have mutations in HA that reduce neutralization by pre-existing antibody immunity (Bedford et al. 2015; Smith et al. 2004; Łuksza and Lässig 2014; Huddleston et al. 2020; Neher et al. 2016; Kikawa et al. 2026b); such mutations increase viral fitness by making a larger fraction of the human population susceptible to infection (Kim et al. 2024; Petrie et al. 2016; Hobson et al. 1972). Therefore, identifying strains and mutations that reduce antibody neutralization is important for vaccine-strain selection. However, viral fitness also depends on inherent transmissibility (which can be affected by mutations in multiple gene segments) (Raghwani et al. 2017; Liu et al. 2024), pre-existing antibody immunity to the other surface protein neuraminidase (Krammer 2019; Monto et al. 2015), and to a lesser extent pre-existing T-cell immunity (Machkovech et al. 2015; Boon et al. 2002). Therefore, measuring how the HAs of different viral strains are neutralized by human antibodies is an important component of viral forecasting—but the strains with the lowest neutralizing titers may not always be the most successful ones.

The last year of evolution of seasonal H3N2 and H1N1 influenza has seen the spread of new viral subclades with reduced neutralization by human HA-directed neutralizing antibodies. In the late summer of 2025, subclades K and D.3.1.1 arose within H3N2 and H1N1 subtypes, respectively, and rapidly increased in frequency. Subclade K has six HA1 mutations relative to its parent subclade J.2.4, mostly in antigenic regions A and B (Liu et al. 2026b). Subclade D.3.1.1 has four HA1 mutations relative to parent subclade D.3.1. These subclades were antigenically advanced relative to parent subclades, with most studies measuring a 1.5- to 2-fold decrease in neutralization by human sera of subclade K (Kikawa et al. 2026a; Kirsebom et al. 2025; Cheng et al. 2026; Dee et al. 2026; Ikonen et al. 2026; Separovic et al. 2026; Wang et al. 2026; Liu et al. 2026b) and a smaller but measurable decrease in neutralization of D.3.1.1 (Kikawa et al. 2026a). Descendant variants of both subclade K and D.3.1.1 are now spreading in the human population, underscoring the continuing need to rapidly measure the antigenic properties of new influenza strains.

Sequencing-based neutralization assays can measure the titers of many sera against many viral strains (Kikawa et al. 2026b; Loes et al. 2024). These assays are higher throughput than conventional neutralization or hemagglutination-inhibition (HAI) assays because they use a sequencing-based readout to measure each serum’s neutralization of >100 viral strains at once. In prior work (Loes et al. 2024) we found that H1N1 sequencing-based neutralization assay titers were similar to those measured in conventional single-cycle neutralization assays, and highly correlated with but quantitatively higher than HAI titers (with a 2-fold change in sequencing-based neutralization assay titer corresponding to a slightly less than 2-fold change in HAI titer). In September of 2025, we started using sequencing-based neutralization assays to make near real-time measurements of the human neutralizing antibody landscape against the current diversity of seasonal H3N2 and H1N1 influenza strains, with these measurements timed to help inform vaccine update decisions (Kikawa et al. 2025; Kikawa et al. 2026a). Here we continue this effort by measuring the neutralization of 148 human H3N2 or H1N1 seasonal strains by 355 recently collected human sera, providing 52,268 total titer measurements that identify new naturally emerging viral strains with HA mutations that reduce recognition by human neutralizing antibody immunity.

## Results

### A library of naturally occurring H3N2 and H1N1 HAs that cover the current diversity of human seasonal influenza

For neutralization titer measurements to be maximally useful for vaccine-strain update decisions, the influenza viruses being assayed should be representative of the current diversity immediately preceding this decision. Towards this end, we designed a library of influenza HAs with the goal of covering the diversity of H3N2 and H1N1 HAs circulating in humans leading up to the September 2026 vaccine-composition meeting.

In late May 2026, we surveyed all human seasonal HAs available in the GenBank or GISAID (Shu and McCauley 2017) databases at that time, and used a combination of computational analysis and manual expert curation to choose a set of HAs from naturally occurring strains based on the following criteria: high overall frequency, rapid increase in frequency over the last six months (Abousamra et al. 2024), mutations at previously established antigenic regions or near/in the receptor binding region (Koel et al. 2013; Wolf et al. 2006; Caton et al. 1982), mutations arising more recurrently than expected by chance (Bloom and Neher 2023; Turner et al. 2026), and observation over a wide geographic range. Overall, this process yielded a set of 77 recently circulating H3N2 HAs and 63 recently circulating H1N1 HAs. We augmented this set with recent egg- and cell-based vaccine strains, for an additional six H1N1 HAs and eight H3N2 HAs. Of the 154 strains we designed for inclusion in the library, 148 could be experimentally generated at sufficient titer, yielding an overall library of 82 H3N2 and 66 H1N1 strains (**Supplementary File 1**). This library largely covers the diversity of recently sequenced human H3N2 and H1N1 influenza, with the HA1 protein sequence of most sequenced human strains over the last year either identical or within one amino-acid mutation of a strain in our library (**Figure 1**).

**Figure 1.**
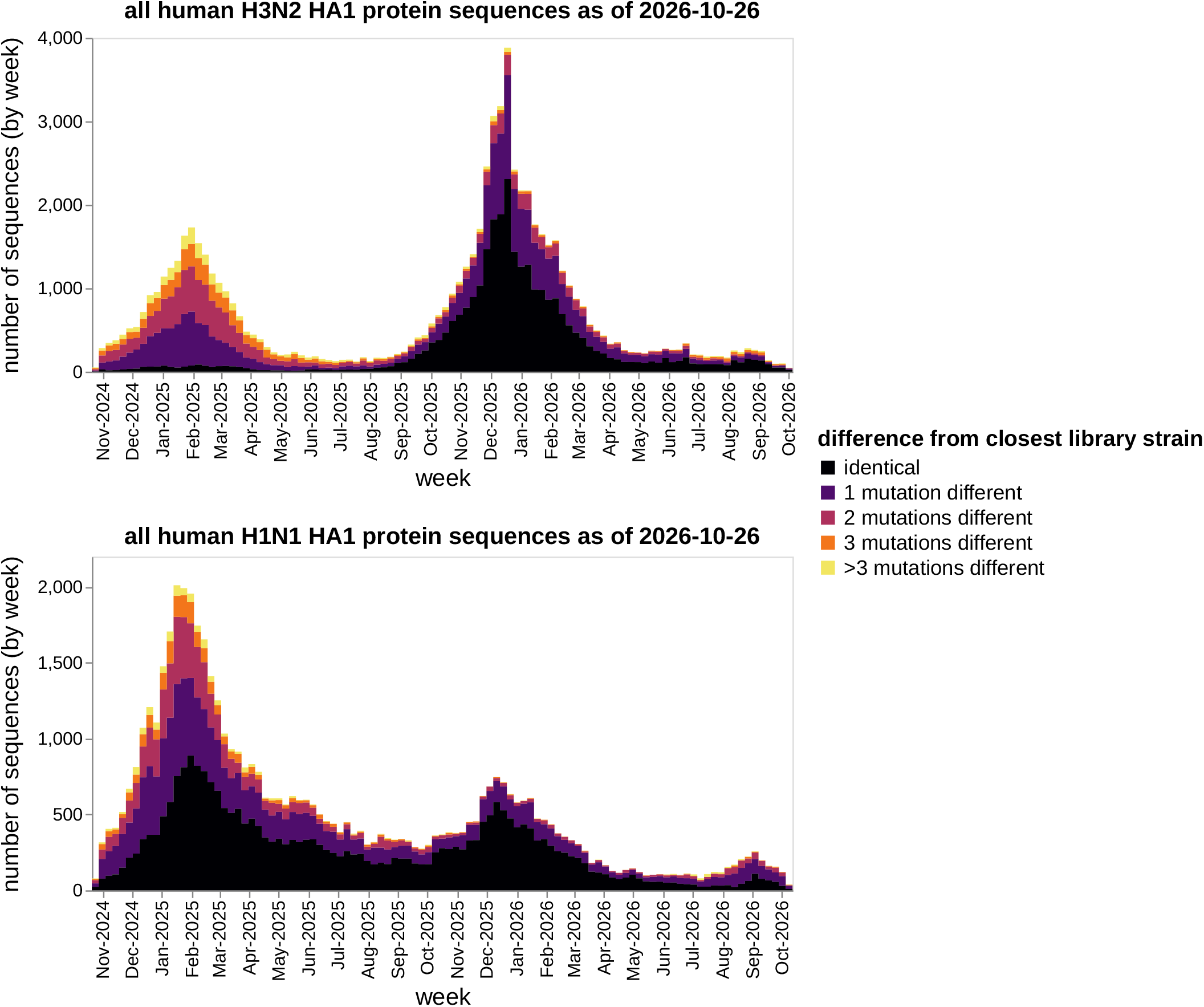
The sequencing-based neutralization assay library largely covers the HA1 diversity of human seasonal H3N2 and H1N1 influenza over the last ~2 years. Number of human seasonal H3N2 (top) or H1N1 (bottom) HA sequences in GISAID (as of October 26, 2026) binned by collection week and colored by HA1 amino-acid identity to the closest strain in our sequencing-based neutralization assay library.

To physically construct this virus library, we individually generated barcoded viruses expressing each HA (Kikawa et al. 2026b; Loes et al. 2024). These viruses contain barcoded HAs with ectodomain protein sequences identical to recently circulating human strains, and all other gene segments from the lab-adapted A/WSN/1933 strain. The viruses were then pooled with the goal of representing all strains at roughly equal titers of transcriptionally active particles, with the pooling done by combining all strains at equal volume, infecting cells, using barcode deep sequencing to quantify the transcriptional titer of each strain, and then re-pooling to balance these titers.

### A set of recently collected human sera

We assembled 355 human sera collected from 260 unique individuals between April and August of 2026 (**Table 1** and **Supplementary File 2**). We included sera from individuals who spanned a wide range of ages (0.5 to 82 years), as past influenza exposure history (which correlates with age) is known to create strong immune imprinting that can affect strain-specific neutralizing titers (Cobey and Hensley 2017). The sera were from three cohorts in the United States, as well as vaccination cohorts in Australia.

**Table 1.**
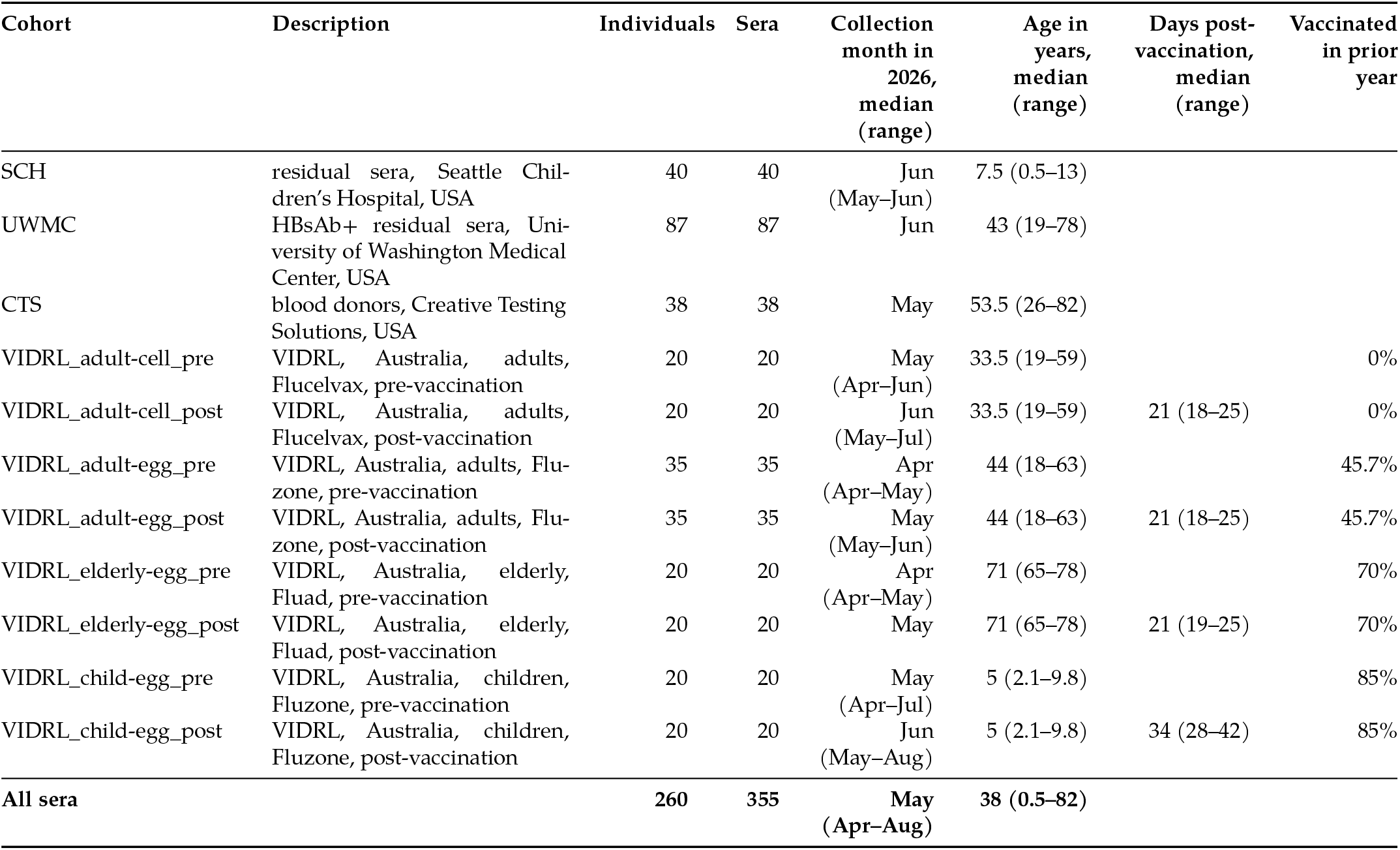
Human sera used in this study. Vaccination status in prior years is not known for the SCH, UWMC, and CTS sera.

| Cohort | Description | Individuals | Sera | Collection month in 2026, median (range) | Age in years, median (range) | Days post-vaccination, median (range) | Vaccinated in prior year |
|---|---|---|---|---|---|---|---|
| SCH | residual sera, Seattle Chil-dren’s Hospital, USA | 40 | 40 | Jun (May–Jun) | 7.5 (0.5–13) |  |  |
| UWMC | HBsAb+ residual sera, Uni-versity of Washington Medical Center, USA | 87 | 87 | Jun | 43 (19–78) |  |  |
| CTS | blood donors, Creative Testing Solutions, USA | 38 | 38 | May | 53.5 (26–82) |  |  |
| VIDRL_adult-cell_pre | VIDRL, Australia, adults, Flucelvax, pre-vaccination | 20 | 20 | May (Apr–Jun) | 33.5 (19–59) |  | 0% |
| VIDRL_adult-cell_post | VIDRL, Australia, adults, Flucelvax, post-vaccination | 20 | 20 | Jun (May–Jul) | 33.5 (19–59) | 21 (18–25) | 0% |
| VIDRL_adult-egg_pre | VIDRL, Australia, adults, Flu-zone, pre-vaccination | 35 | 35 | Apr (Apr–May) | 44 (18–63) |  | 45.7% |
| VIDRL_adult-egg_post | VIDRL, Australia, adults, Flu-zone, post-vaccination | 35 | 35 | May (May–Jun) | 44 (18–63) | 21 (18–25) | 45.7% |
| VIDRL_elderly-egg_pre | VIDRL, Australia, elderly, Fluad, pre-vaccination | 20 | 20 | Apr (Apr–May) | 71 (65–78) |  | 70% |
| VIDRL_elderly-egg_post | VIDRL, Australia, elderly, Fluad, post-vaccination | 20 | 20 | May | 71 (65–78) | 21 (19–25) | 70% |
| VIDRL_child-egg_pre | VIDRL, Australia, children, Fluzone, pre-vaccination | 20 | 20 | May (Apr–Jul) | 5 (2.1–9.8) |  | 85% |
| VIDRL_child-egg_post | VIDRL, Australia, children, Fluzone, post-vaccination | 20 | 20 | Jun (May–Aug) | 5 (2.1–9.8) | 34 (28–42) | 85% |
| **All sera** |  | **260** | **355** | **May (Apr–Aug)** | **38 (0.5–82)** |  |  |

The sera from the United States included remnant pediatric sera from hospital visits (mostly unrelated to viral infections) to Seattle Children’s Hospital (SCH cohort), remnant adult sera from routine hepatitis B surface antibody testing at the University of Washington Medical Center in Seattle (UWMC cohort), and sera from blood donors across the United States from Creative Testing Solutions (CTS cohort).

The sera from Australia were provided by the Victorian Infectious Diseases Reference Laboratory (VIDRL), and were collected from children, adults, and elderly individuals. These individuals were sampled pre- and post-vaccination with a 2026 Southern Hemisphere cell-based (Flucelvax) or egg-based (Fluzone or Fluad) seasonal influenza vaccine, which contain a J.2.4 H3N2 strain and a D.3.1 H1N1 strain for the cell-based vaccine, and a J.2.4:F195Y strain and D.3.1:Q223R strain for the egg-based vaccine. As detailed in **Table 1**, there are differences between the groups that received the different vaccines in terms of post-vaccination sampling time and prior vaccination history.

### Some H3N2 subclade K variants have reduced neutralization, especially those with mutations in antigenic region D

The neutralization titers against recent H3N2 strains are shown in **Figure 2** and reported in **Supplementary File 3**; see also https://jbloomlab.github.io/flu-seqneut-2026/summary.html for interactive plots that are easier to explore than the static figures in this manuscript.

**Figure 2.**
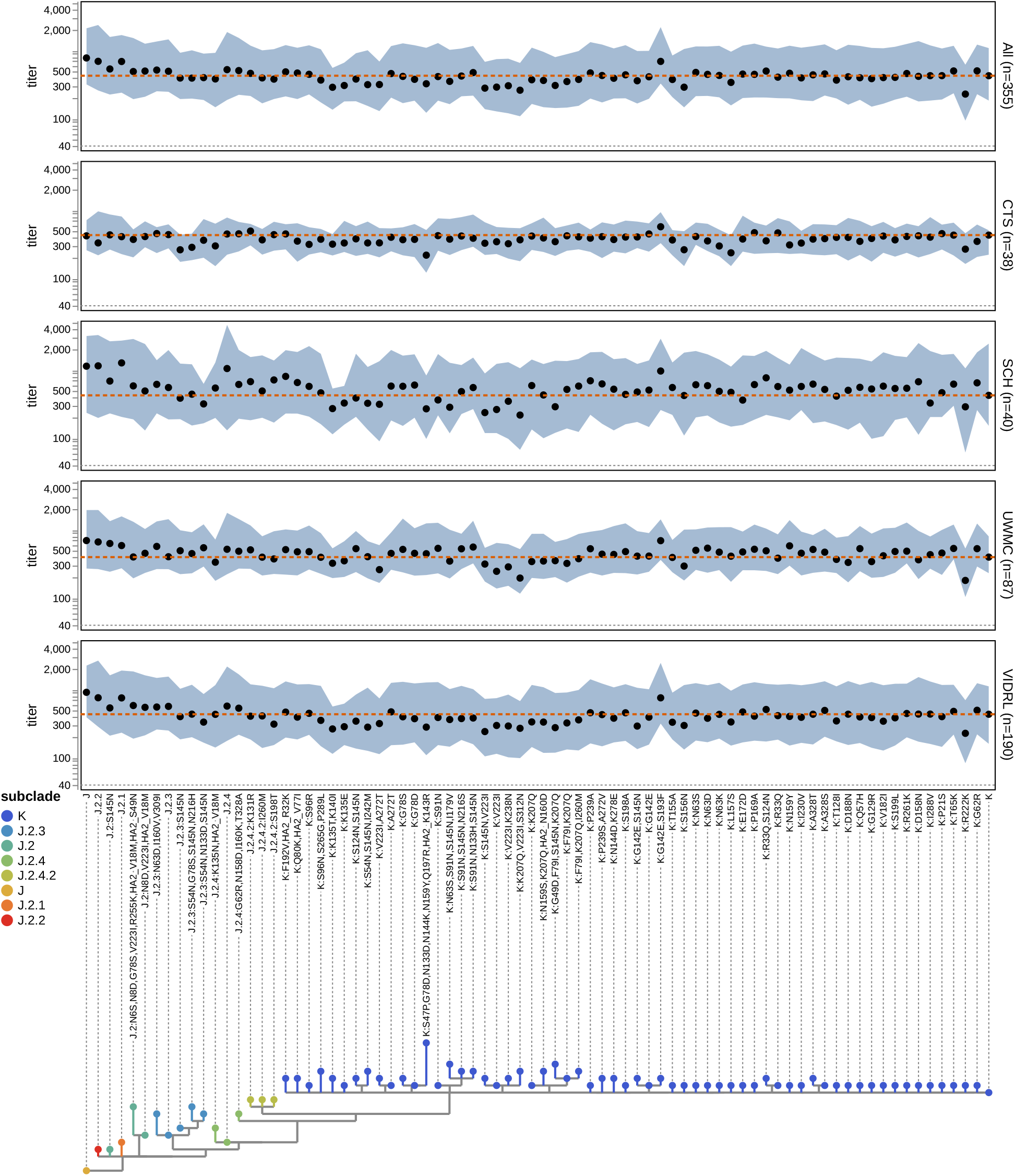
Neutralization titers of all tested sera against recent H3N2 HAs. Points indicate median titer and shaded regions indicate interquartile range. The dashed orange line indicates the titer against subclade K (the 2026-2027 cell-based vaccine strain) and the dotted gray line indicates the lower limit of detection (a titer of 40). Labels to the right of each plot indicate the sera set and number of sera in that set. X-axis labels indicate the HA protein haplotype (subclade and any derived mutations), and the phylogenetic tree shows the relationships among the protein sequences and is colored by subclade.

Across all sera, the lowest H3N2 titers are to subclade K and K-descendant strains. Strains from other recent subclades (J, J.2, J.2.1, J.2.2, J.2.3, and J.2.4) are neutralized by recent human sera similarly or more potently than the parental subclade K strain that is the H3N2 component of the 2026-2027 Northern Hemisphere vaccine (these strains are mostly at or above the orange dashed line in **Figure 2**). Notably, these other non-K H3N2 sub-clades have greatly decreased in frequency in the human population over the last nine months, coincident with the rapid spread of subclade K.

Within subclade K, there are new strains with additional HA mutations that are neutralized substantially less well than the parental subclade K (below the orange dashed line in **Figure 2**). The subclade K strains with the lowest neutralization mostly carry mutations at HA sites 223 or 222 (e.g., V223I or R222K). Both these sites are in antigenic region D, which was not mutated in the parental subclade K strain—which instead had mutations in antigenic regions A and B relative to its parent subclade J.2.4 (**Figure 3**). Therefore, mutations in antigenic region D may impair recognition by the antibodies that could still cross-neutralize both subclade K and older strains (Liu et al. 2026a; Kikawa et al. 2026a). Notably, H3N2 HA had an identity of I223 from ~2010 until the emergence of subclade J in ~2022, so some individuals may have antibodies from prior exposures that target I223 variants although overall we still observe V223I to typically cause a decrease in neutralization in subclade K. A mutation at site 222 has only fixed once in human H3N2 evolution (W222R in ~2000). Subclade K strains with mutations at other antigenic region A and B sites (e.g., 145, 156, 157) also had reduced neutralization by some sera relative to the parental subclade K (**Figure 2**). See https://nextstrain.org/community/jbloomlab/flu-seqneut-2026@main/H3N2 for an interactive Nextstrain tree that visualizes the titers alongside all mutations.

**Figure 3.**
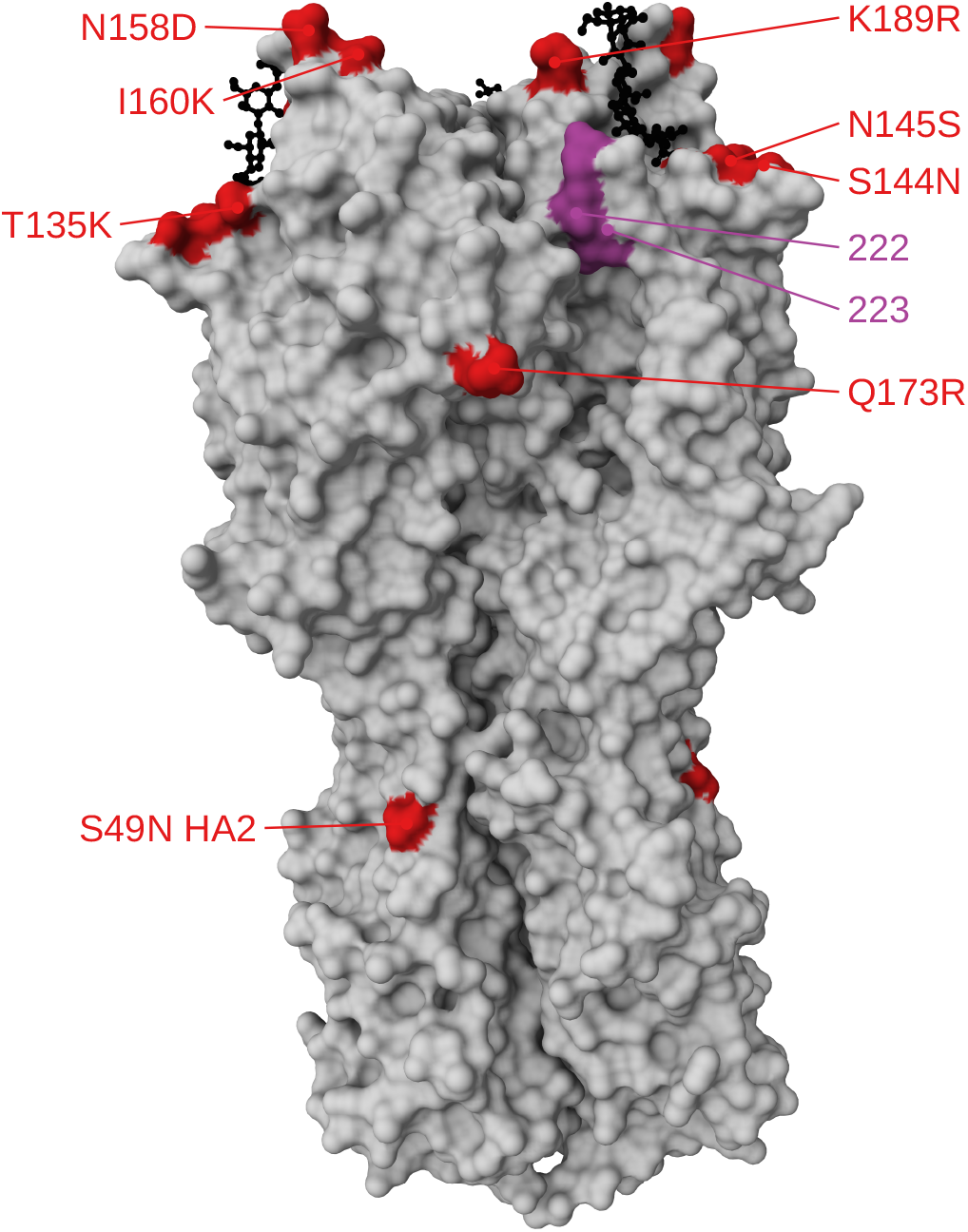
Structure of H3 HA indicating sites 222 and 223 and mutations that separate the 2025-2026 cell-based vaccine (J.2:S145N) and subclade K. Structure of H3 trimer (PDB 8faw) with red indicating mutations that separate the 2025-2026 cell-based vaccine from subclade K, and purple indicating sites 222 and 223 where mutations to subclade K further reduce neutralization. A sialic-acid receptor analogue is shown in black sticks. Sites 135, 144, and 145 are in antigenic region A; sites 158, 160, and 189 are in antigenic region B; sites 222 and 223 are in antigenic region D.

A subclade K strain with mutations G142E and S193F was more potently neutralized by most sera than the parental subclade K strain (**Figure 2**). This increased neutralization by most sera is likely because S193F reverts this antigenic region B site to its amino-acid identity prior to ~2020; consistent with this hypothesis, the K:G142E,S193F strain is neutralized similarly to other subclade K strains by sera from young children (less than five years of age) who would not have been exposed to F193-carrying viruses or vaccines (see the interactive version of **Figure 2** at https://jbloomlab.github.io/flu-seqneut-2026/summary.html that allows subsetting of sera by subject age using the sliders below the plot).

### H1N1 subclade D.3.1.1 has reduced neutralization, with Sa mutations strongly affecting a subset of sera

The neutralization titers against recent H1N1 strains are shown in **Figure 4** and reported in **Supplementary File 3**; see also the summary website for interactive plots.

**Figure 4.**
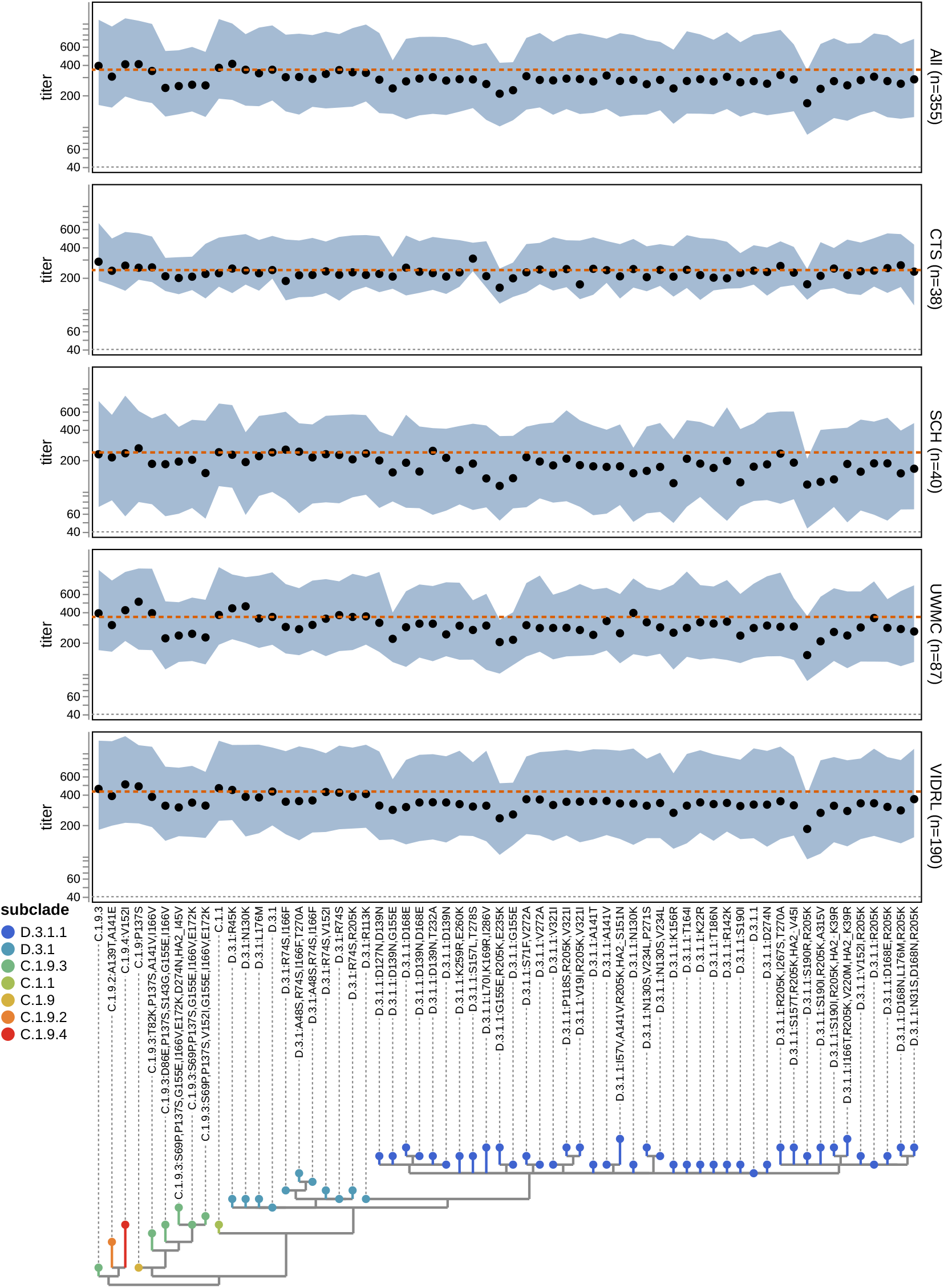
Neutralization titers of all tested sera against recent H1N1 HAs. Points indicate median titer and shaded regions indicate interquartile range. The dashed orange line indicates the titer against subclade D.3.1 (the 2026 and 2026-2027 cell-based vaccine strain), and the dotted gray line indicates the lower limit of detection (a titer of 40). Labels to the right of each plot indicate the sera set and number of sera in that set. X-axis labels indicate the HA protein haplotype (subclade and any derived mutations), and the phylogenetic tree shows the relationships among the protein sequences and is colored by subclade.

The 2026 Southern Hemisphere and 2026-2027 Northern Hemisphere H1N1 vaccine strain is a subclade D.3.1 strain, but nearly all recent human H1N1 sequences are from the D.3.1-descended subclade D.3.1.1. Most D.3.1.1 strains are neutralized more poorly by recent human sera than D.3.1 (below the dashed orange line in **Figure 4**). A handful of highly mutated strains descended from sub-clade C.1.9.3 are also poorly neutralized (**Figure 4**), although C.1.9.3 strains have not recently been observed in actual sequencing of human H1N1 influenza.

The lowest titers are to new D.3.1.1 strains with additional HA mutations (**Figure 4**). The strain with the lowest overall titers is D.3.1.1:S190R,R205K. Strains with G155E and to a lesser extent S157L (both in antigenic region Sa) also had clearly reduced titers. See https://nextstrain.org/community/jbloomlab/flu-seqneut-2026@main/H1N1 for an interactive Nextstrain tree that visualizes the titers alongside all mutations.

While strains with G155E are neutralized less well across all sera (**Figure 4**), the pattern is particularly striking if the sliders are used to subset on sera from individuals ages 15 to 25 years in the inter-active plots at https://jbloomlab.github.io/flu-seqneut-2026/summary.html. The same trend can be seen in **Figure 5**, which stratifies sera based on whether or not D.3.1.1:G155E is neutralized at least two-fold worse than D.3.1.1. Sera that are strongly affected by G155E tend to be from teenagers and young adults and have higher overall H1N1 titers, but dramatically decreased titers to G155E-containing strains (**Figure 5**). A similar trend occurs to a lesser extent for S157L. Sites 155 and 157 are adjacent in antigenic region Sa at a structural location distinct from the other mutations that separate subclade D.3.1.1 from its parent subclade D.3.1 (**Figure 6**). Interestingly, G155E was described shortly after the 2009 H1N1 pandemic as a mutation that greatly reduced neutralization by sera from ferrets immunized with an early 2009 H1N1 pandemic strain but had little impact on sera from humans imprinted with pre-2009 H1N1 (Li et al. 2013; Chen et al. 2010, note G155E is G158E in H3 numbering). The age group in which G155E has the largest effect in our data is humans who were likely first exposed to H1N1 during or shortly after the 2009 pandemic, whereas the older individuals in which it has less effect would have been imprinted with the older H1N1 strains that preceded the 2009 pandemic. Notably, G155E has recently spread at high frequency in multiple D.3.1.1 H1N1 sub-lineages after previously remaining rare among human H1N1 influenza since 2009.

**Figure 5.**
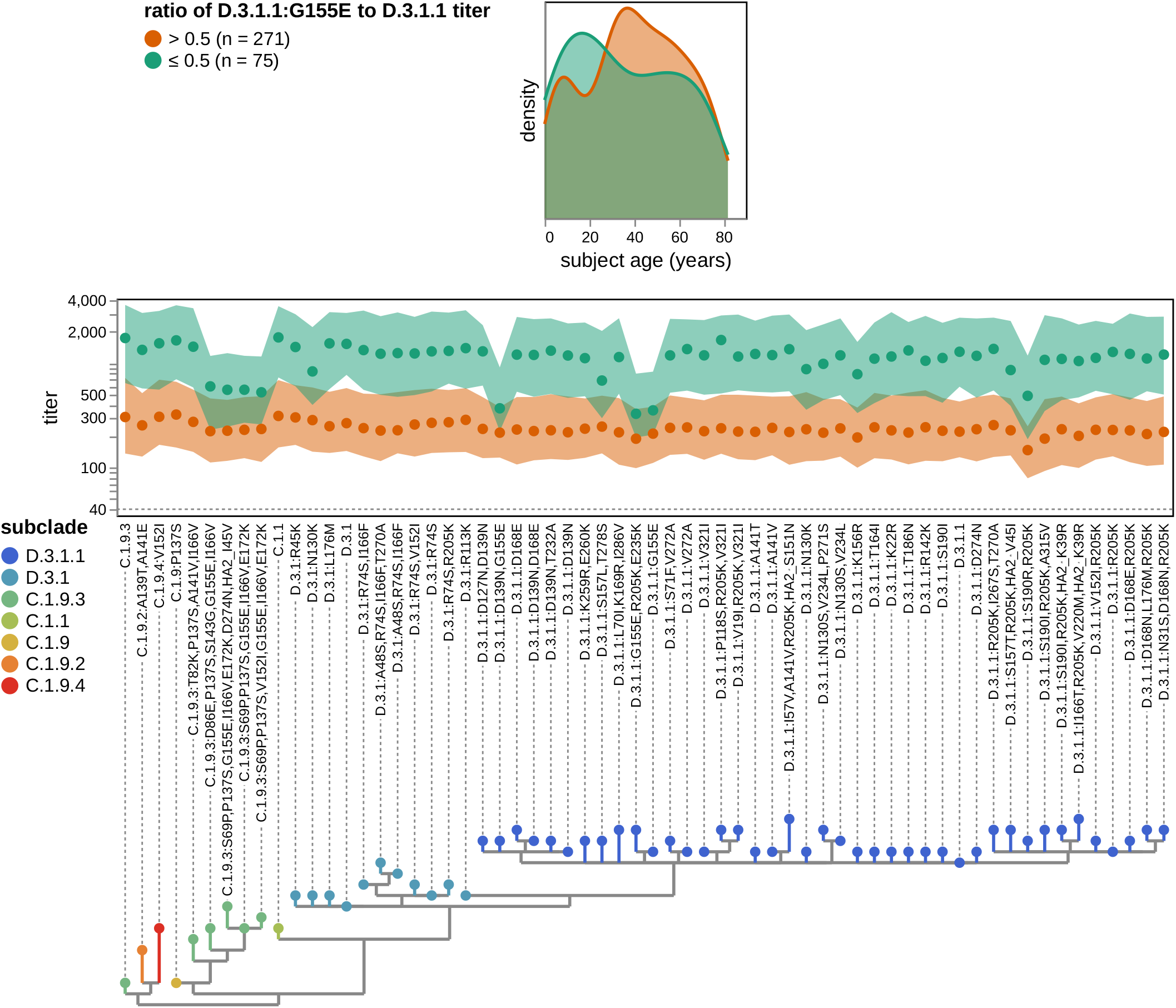
G155E in H1N1 HA causes a large decrease in neutralization for a subset of individuals who otherwise have high titers and tend to be teenagers or young adults. All sera stratified by whether the G155E mutation in D.3.1.1 reduces neutralization titers by more than two-fold relative to D.3.1.1 without any additional mutations. The top density plot shows the age distributions for both groups, and the lower plot shows the median (points) and interquartile range (shaded region) neutralization titers against recent H1N1 strains. In both plots, green indicates the 75 sera for which G155E causes a more than two-fold titer drop, and orange indicates the 271 sera for which G155E does not cause such a large titer drop. The dotted gray line in the titer plot indicates the lower limit of detection (a titer of 40). The handful of sera for which the titer against either D.3.1.1 or D.3.1.1:G155E was at the lower limit of detection are not shown in this plot to avoid confounding effects due to truncated measurements.

**Figure 6.**
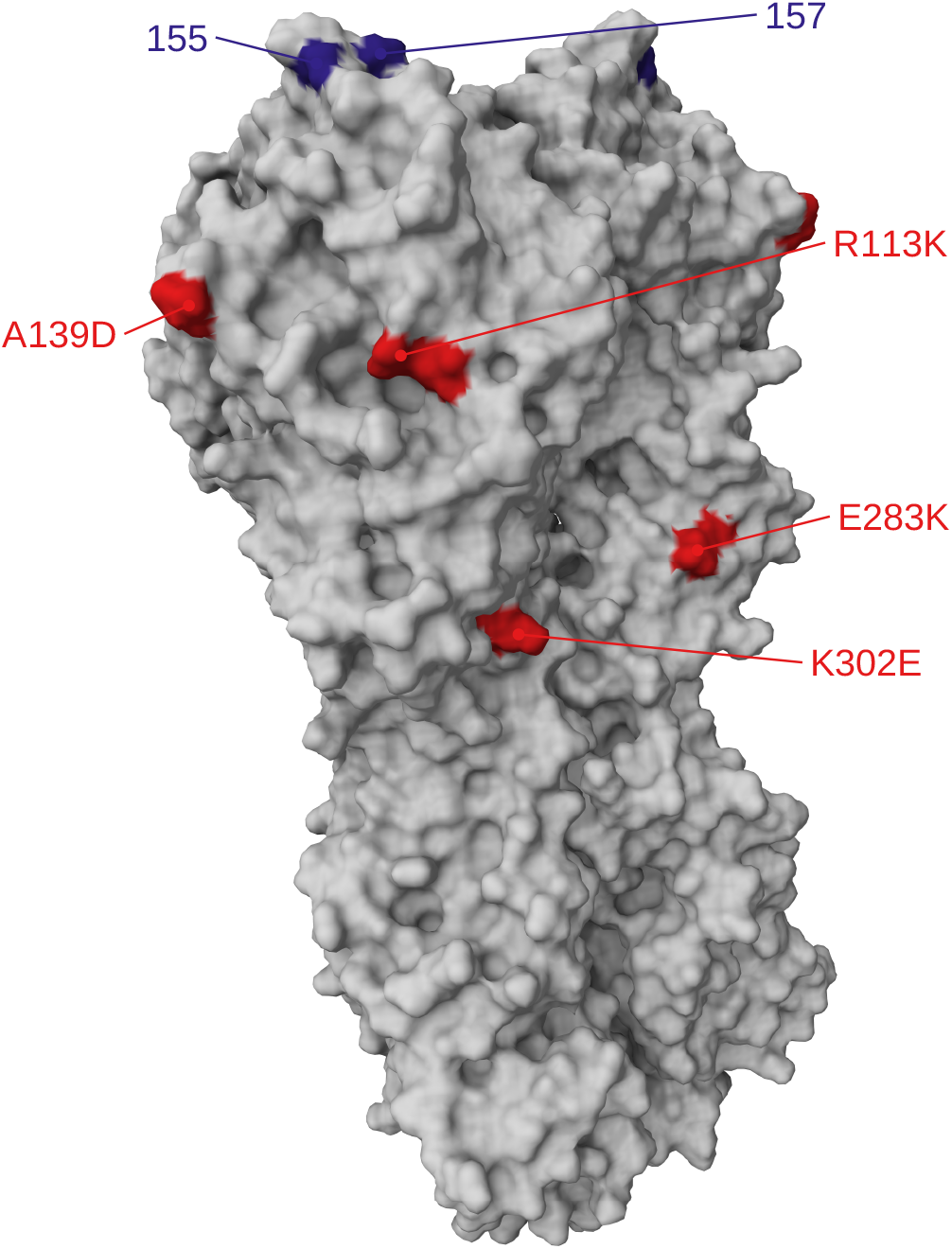
Structure of H1 HA indicating sites 155 and 157 and mutations that separate the 2026 and 2026-2027 cell-based vaccine (D.3.1) and subclade D.3.1.1. Structure of H1 trimer (PDB 9gsp) with red indicating mutations that separate D.3.1 from D.3.1.1, and blue indicating sites 155 and 157 where new mutations in D.3.1.1 further reduce neutralization. Site 139 is in antigenic region Ca2; sites 155 and 157 are in antigenic region Sa.

### The 2026 Southern Hemisphere vaccine boosts titers to all current H3N2 and H1N1 strains, with modest strain-to-strain variation

The Australia-based VIDRL sera were from four groups sampled before and after administration of the 2026 Southern Hemisphere influenza vaccine: children who received the egg-based Fluzone vaccine, adults who received the egg-based Fluzone vaccine, adults who received the cell-based Flucelvax vaccine, and elderly individuals who received the adjuvanted egg-based Fluad vaccine (**Table 1** and **Supplementary File 2**). In addition to the differences in age and vaccine formulations, these groups differ in two other factors that can affect vaccine response: the timing of the post-vaccination serum collection (Lane et al. 2025) and extent to which they had been vaccinated in prior years (Cowling et al. 2024) (**Table 1**). Therefore, we show the adult Fluzone co-hort stratified by prior-year vaccination status. Even so, we focus on trends in the strain-specific response within groups rather than on differences in the absolute titer increase after vaccination across groups, since the latter property could be affected by multiple factors.

Vaccination increased titers to all recent strains for both H3N2 and H1N1, so the post-vaccination strain-specific titers largely conform to the overall trends described in the preceding two sections (see **Figure 7** and **Figure 8** for raw titers, and the summary website for interactive plots that also show the fold change in titer after vaccination). The H3N2 strain in the 2026 Southern Hemisphere vaccine is J.2.4 for the cell-based vaccine and J.2.4:F195Y for the egg-based vaccine, and the H1N1 strain is D.3.1 for the cell-based vaccine and D.3.1:Q223R for the egg-based vaccine. However, vaccination did not induce a notably larger increase in titers against the matched J.2.4 and D.3.1 strains compared to more antigenically advanced subclade K and D.3.1.1 strains. Instead, the strain-specific post-vaccination titers mostly mirrored the pre-vaccination titers but generally shifted higher (**Figure 7** and **Figure 8**). For H3N2 there were no strains to which vaccination consistently induced a lower titer boost across groups: for instance, the adult Fluzone group with a prior-year vaccination had a low titer boost to K:K135E, but other groups (including adults who received Flucelvax, children who received Fluzone, and adults receiving Fluzone without a prior-year vaccination) had a similar titer increase to the K:K135E as to other strains. For H1N1 there was a discernible trend for vaccination to induce a slightly lower titer boost to some strains (particularly those containing G155E), but the magnitude of this trend was modest (**Figure 9**).

**Figure 7.**
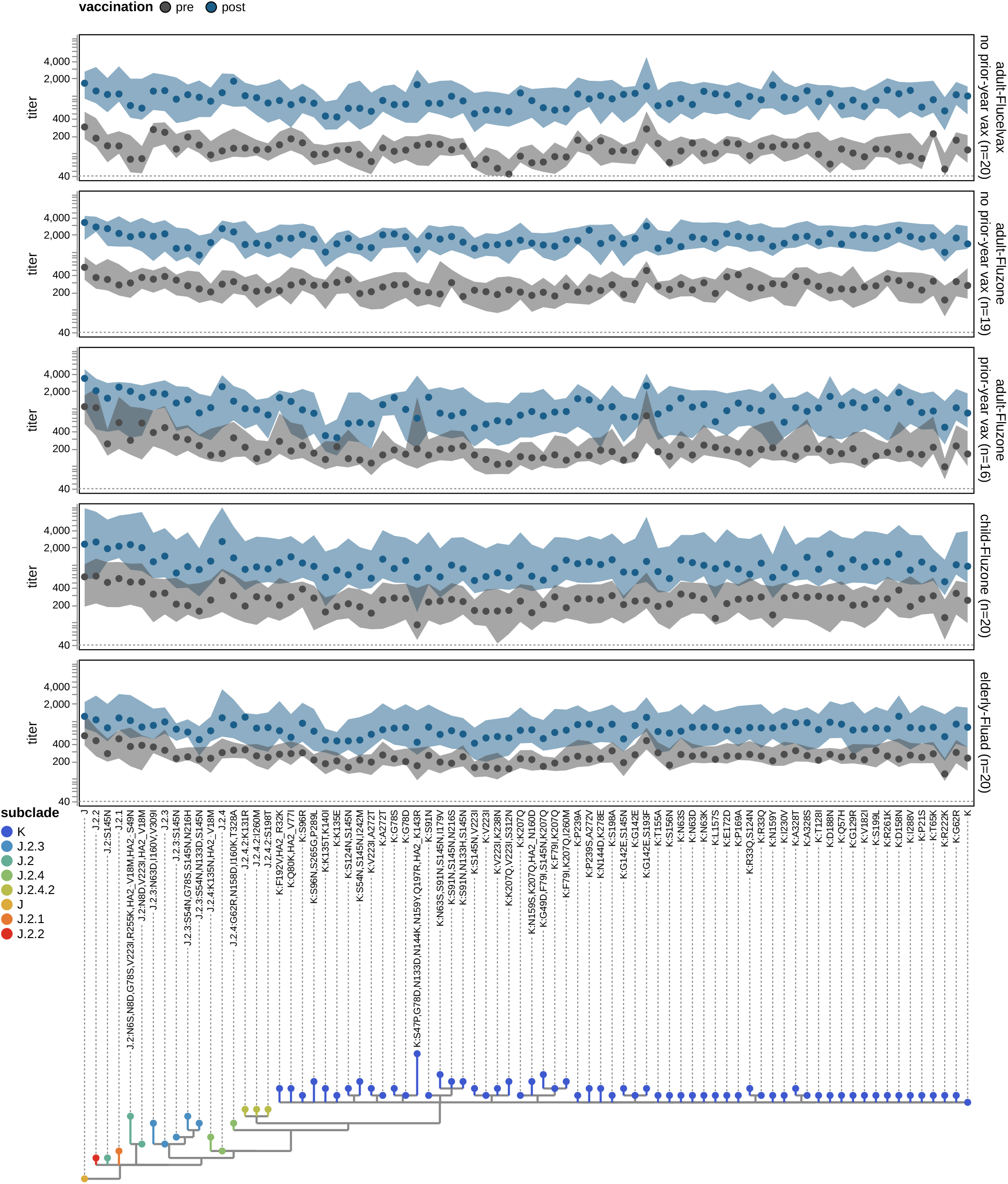
Neutralization titers pre- and post-vaccination against recent H3N2 HAs. Each plot shows the titers for a different cohort from the VIDRL (Australia) sera set before and after vaccination with the 2026 Southern Hemisphere vaccine (J.2.4 for the cell-based Flucelvax vaccine, and J.2.4:F195Y for the egg-based Fluzone and Fluad vaccines). The adult Fluzone cohort is further stratified by prior-year vaccination status; all adult Flucelvax recipients were not vaccinated in the prior year. Points indicate the median and shaded regions indicate the interquartile range, with gray indicating pre-vaccination titers and blue indicating post-vaccination titers. The dotted gray line indicates the lower limit of detection (a titer of 40). Post-vaccination sera were collected a median of three weeks after vaccination for the adult and elderly cohorts and five weeks after vaccination for the child sera; see **Table 1**.

**Figure 8.**
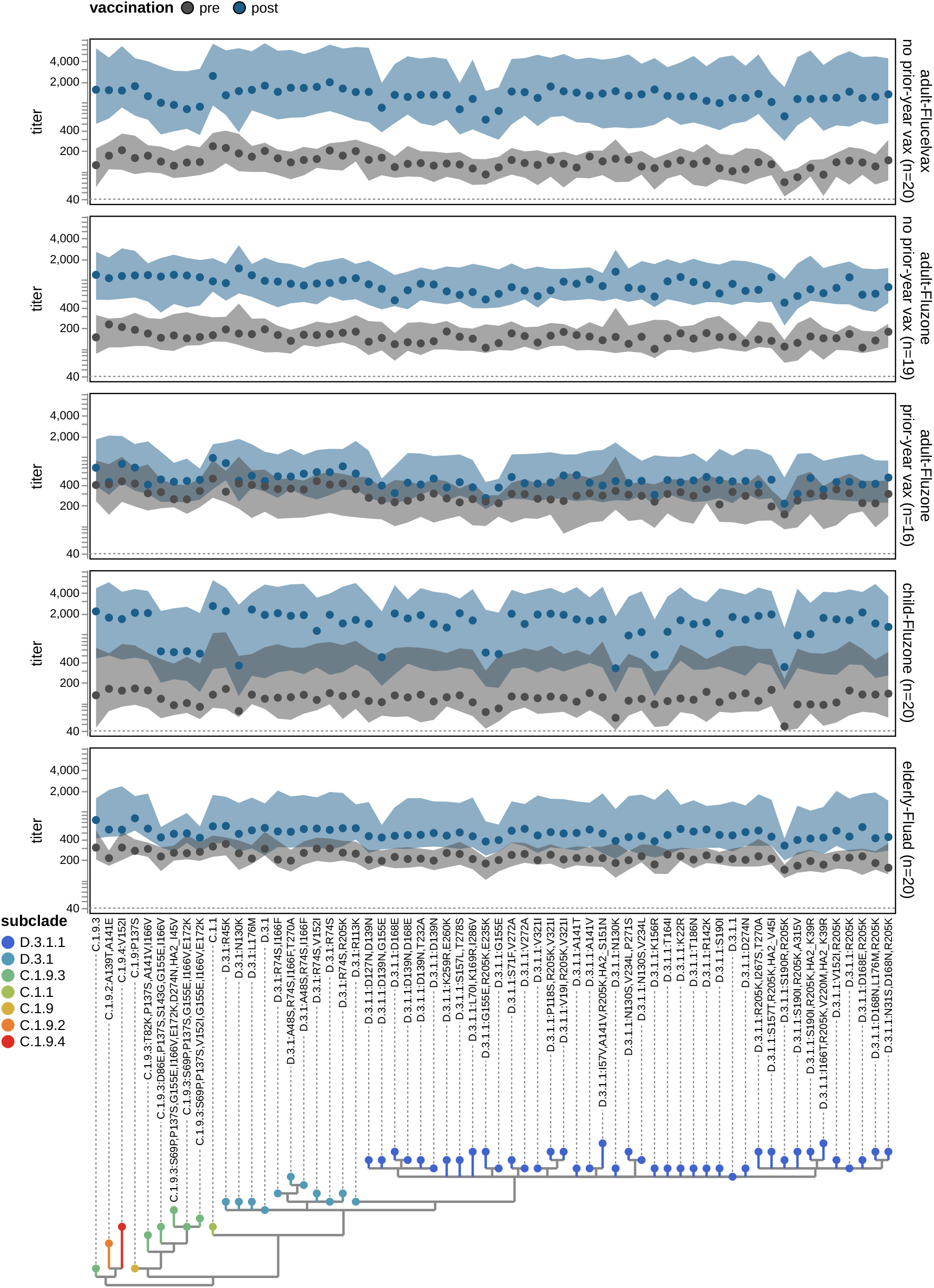
Neutralization titers pre- and post-vaccination against recent H1N1 HAs. Each plot shows the titers for a different cohort from the VIDRL (Australia) sera set before and after vaccination with the 2026 Southern Hemisphere vaccine (D.3.1 for the cell-based Flucelvax vaccine, and D.3.1:Q223R for the egg-based Fluzone and Fluad vaccines). The adult Fluzone cohort is further stratified by prior-year vaccination status; all adult Flucelvax recipients were not vaccinated in the prior year. Points indicate the median and shaded regions indicate the interquartile range, with gray indicating pre-vaccination titers and blue indicating post-vaccination titers. The dotted gray line indicates the lower limit of detection (a titer of 40). Post-vaccination sera were collected a median of three weeks after vaccination for the adult and elderly cohorts and five weeks after vaccination for the child sera; see **Table 1**.

**Figure 9.**
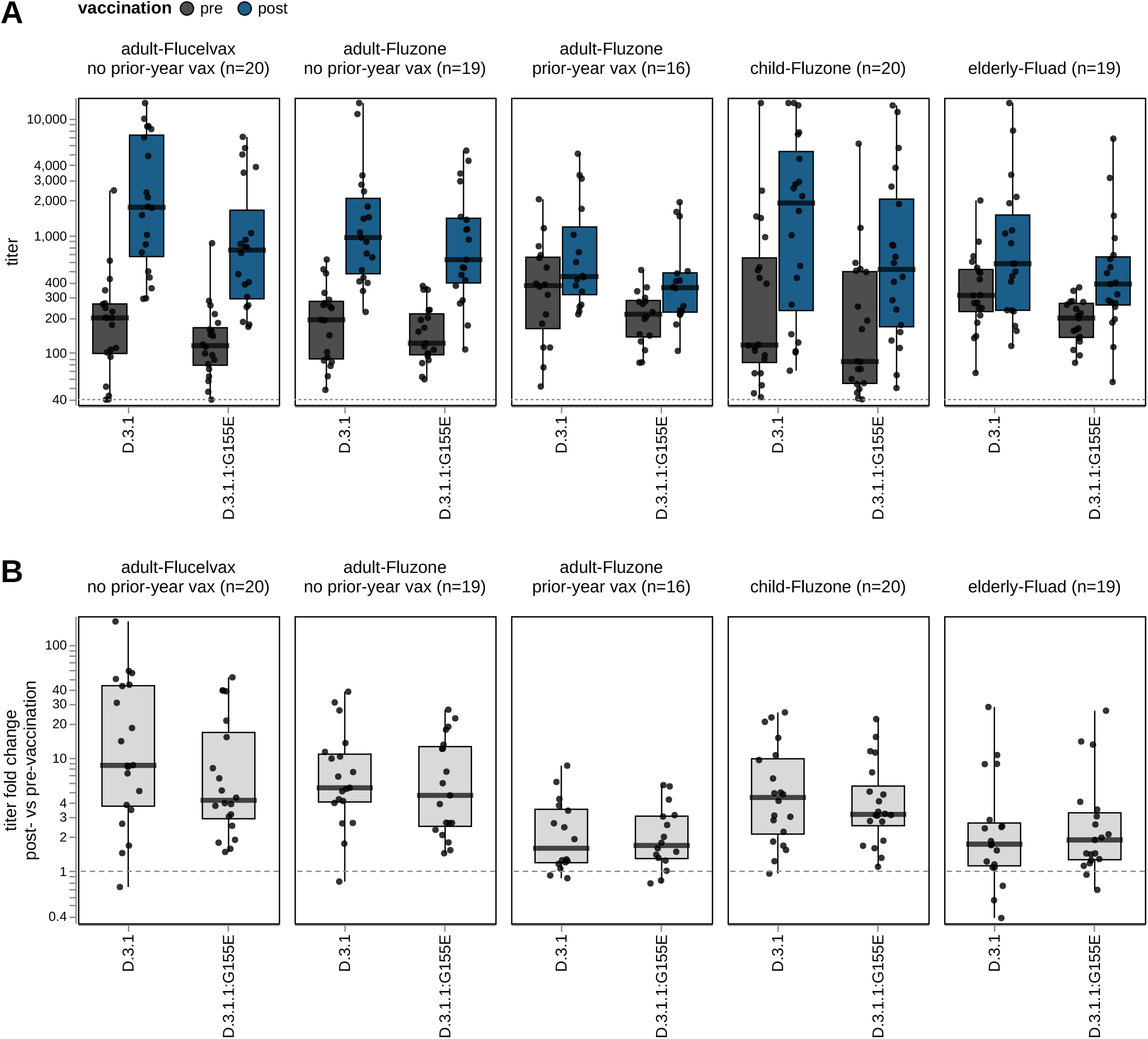
Changes in neutralization titers after vaccination for the cell-based vaccine strain D.3.1 and the D.3.1.1:G155E strain. Distributions of (**A**) neutralization titers (pre-vaccination in dark gray and post-vaccination in blue) and (**B**) titer fold changes against D.3.1 (the cell-based vaccine strain) and D.3.1.1:G155E. The boxes show the interquartile range, the thick black lines indicate the median, and the whiskers show the full range. In panel A, each point represents an individual at one timepoint, and in panel B, each point represents the fold change of post-vaccination titer over pre-vaccination titer. Only individuals with both pre- and post-vaccination titers against both strains are shown. The dotted gray lines represent the lower limit of detection (a titer of 40) in panel **A** and no change between pre- and post-vaccination titers in panel **B**.

## Discussion

We have measured how a large set of human sera neutralize current human H3N2 and H1N1 influenza strains. Our results show that the two subclades that spread widely over the last year (subclade K for H3N2 and D.3.1.1 for H1N1) are now spawning descendants with reduced neutralization by human sera. These descendant strains have HA mutations that are associated with decreased neutralization, some of which have arisen independently in different combinations. For H3N2, mutations at sites in antigenic region D (e.g., 223 and 222) have arisen recurrently and reduce neutralization; mutations at sites in antigenic regions A and B are also present in strains with reduced neutralization (e.g., 145, 156, 157). For H1N1, strains with mutations at sites in antigenic region Sa (e.g., 155 and 157) have arisen recurrently and reduce neutralization; mutations at sites in antigenic regions Sb and Ca1 (e.g., 190 and 205) are also present in strains with reduced neutralization.

However, the diversity of new antigenic variant strains makes it challenging to use our data alone to determine which strains will dominate a year from now. For H1N1, our data show that D.3.1.1 is antigenically advanced over D.3.1, and virtually all human H1N1 influenza observed over the last few months is D.3.1.1 or a descendant strain.

However, it is uncertain which more antigenically advanced variant of H1N1 will spread over the next year— G155E reduces neutralization of all H1N1 strains and has recently arisen recurrently, but it occurs in multiple genetic backgrounds and there are also other antigenic mutations that could outcompete it. Similarly, although our data identify multiple new antigenic variant strains of subclade K, it remains unclear which of these new variants will dominate over the next year. Ultimately answering this question will require integrating neutralization data with forecasting models that also draw on other data sources (Łuksza and Lässig 2014; Huddleston et al. 2020; Shi et al. 2025; Neher et al. 2016; Chen et al. 2026)— such as sequencing counts (Abousamra et al. 2024) and mutation recurrence (Turner et al. 2026; Bloom and Neher 2023)—to estimate which current strains are most likely to spread in the human population.

Another finding is that despite clear antigenic variation among recent strains, the current 2026 Southern Hemisphere vaccine induced mostly similar increases in neutralization titer to all strains—such that the strain-to-strain variation in post-vaccination titers largely mirrored the variation in pre-vaccination titers. This raises the question of whether a vaccine update is merited even if it is possible to forecast which new strain will spread in the next year. However, we only measured neutralization by serum collected three to five weeks after vaccination when the boosted serum antibodies in individuals with a prior influenza exposure derive largely from a recall response of pre-existing memory B cells (Wrammert et al. 2008; Andrews et al. 2015; Ellebedy et al. 2016)—but influenza vaccines can also recruit naive strain-specific B-cells into germinal centers or lead to affinity maturation (Ellebedy et al. 2020; Turner et al. 2020; Matz et al. 2026; McIntire et al. 2024; Hoehn et al. 2021), a process that might not lead to an immediate change in serum titers. Studies of COVID-19 vaccination found that while a single updated booster vaccination of previously exposed individuals increased titers to both older and newer strains, over multiple years updated boosters partially overcame immune imprinting to preferentially increase titers to new viral variants (Yisimayi et al. 2024; Mellis et al. 2026; Tortorici et al. 2026; Alsoussi et al. 2023). So an updated vaccine can in principle lead to changes in the B-cell compartment that do not strongly manifest at the serological level until later exposures (Schiepers et al. 2023; Yisimayi et al. 2024), although it is unclear if this is actually the case for current influenza vaccines (Davis et al. 2020; Spangler et al. 2025).

Overall, our results underscore the complex hetero-geneity of the human neutralizing antibody landscape (Cobey and Hensley 2017; Lee et al. 2019; Kikawa et al. 2026b). For instance, G155E in H1N1 appears to only modestly decrease neutralization if one simply examines the median across all sera—but in fact this mutation greatly decreases neutralization for some sera but only mildly impacts other sera. Here we have summarized a rich dataset full of such person-to-person heterogeneity as best as possible on the near real-time schedule required to make it useful for pending vaccine update decisions. But an important area for future work is analyzing these data as well as similar data from earlier timeframes (Kikawa et al. 2025; Kikawa et al. 2026a) to better understand how heterogeneous human neutralization data can inform forecasts of viral antigenic evolution.

## Acknowledgements

We thank Michael Lassig, Marta Luksza, Denis Ruchnewitz, Sarah James, and Nicola Lewis for suggestions of which viral strains to include in the sequencing-based neutralization assays. We thank Richard Neher for helpful comments on the data. We thank Dolores Covarrubias, Catherina Artikis for assistance with deep sequencing at the Fred Hutch Genomics Core. We gratefully acknowledge the authors, originating and submitting laboratories of the sequences from the GISAID EpiFlu Database, which were used to inform some of the analyses for this study.

The experimental work and data analysis at the Fred Hutch Cancer Center was supported by the NIH/NIAID under grant R01AI165821 (to JDB and JH), grant F30AI186284 (to CK), and contract 75N93021C00015 (to SEH and JDB). This research utilized the Genomics & Bioinformatics Shared Resource (RRID:SCR_022606) of the Fred Hutch/University of Washington Cancer Consortium (P30 CA015704) and the Fred Hutch Scientific Computing Cluster (NIH grants S10-OD-020069 and S10-OD-028685). The Australian adult and elderly sera were collected for studies led by HSM and SW in part with funding from Seqirus Limited, and the Australian child sera were collected for a study led by ST in part with funding from AstraZeneca; note that this funding was not tied to the current study, but rather the already-collected de-identified sera reagents were additionally shared for the work described here. CK is supported in part by a Beyond the Journal Award from the Experiment Foundation/Navigation Fund. AB is a Coefficient Giving Fellow of the Life Sciences Research Foundation. JDB is an Investigator of the Howard Hughes Medical Institute.

## Author Contributions

CK, AB, JH, SEH, and JDB designed the study. CK and AB performed the experiments. CK, AB, JH, and JDB performed the primary data analysis and wrote the first draft of the manuscript. SAT and DJS shared data used to help choose which viral strains to include. JAE and KL coordinated collection of the Seattle Children’s Hospital sera. MB, ML, MS, and BS coordinated collection of the Creative Testing Solutions sera. ALG provided the University of Washington Medical Center sera. HP and IGB coordinated sharing of the Australian sera. SW and HSM led collection of the Australian adult and elderly sera under a separate pre-existing study, and ST led collection of the Australian child sera under a separate pre-existing study; these sera were then shared for this study. All authors reviewed the final manuscript.

## Methods

### Data and code availability

All data and computer code are at https://github.com/jbloomlab/flu-seqneut-2026. See https://jbloomlab.github.io/flu-seqneut-2026/summary.html for a summary of the results with interactive plots and links to the underlying data, and https://jbloomlab.github.io/flu-seqneut-2026 for more extensive documentation of the neutralization data at a per-plate and per-serum level. The final QC-ed titer data reported in this manuscript are in the CSV files at https://github.com/jbloomlab/flu-seqneut-2026/tree/main/results/final_titer_data and also in **Supplementary File 1, Supplementary File 2**, and **Supplementary File 3**.

### Biosafety

All experiments used viruses with HA protein ectodomains identical to recently circulating human seasonal H3N2 and H1N1 strains available in public databases (GenBank or GISAID) or recent cell- or egg-based vaccine strains. None of the HA ectodomains contained any additional protein mutations relative to recent human strains or vaccine strains. The rest of the HA gene segment and the seven non-HA genes were derived from the lab-adapted A/WSN/1933 (H1N1) strain. Since both recent human seasonal strains and the lab-adapted A/WSN/1933 strain are classified as appropriate for study at biosafety-level 2 by the BMBL handbook (edition 6), this work was approved by the Fred Hutch institutional biosafety committee at biosafety-level 2.

### HA numbering and definition of antigenic regions

Throughout this manuscript we use H3 numbering for the H3N2 HAs and H1 numbering for the H1N1 HAs (these numbering schemes start with 1 for the first residue of the ectodomain of that subtype, and number HA1 and HA2 separately). The antigenic region definitions are taken from (Stray and Pittman 2012) for H3N2 (see https://jbloomlab.github.io/flu-seqneut-2026/H3_prot_struct_viz.html#view=antigenic-regions for these regions projected on the HA structure and their constituent sites defined), and from (Wilson et al. 2015) for H1N1 (see https://jbloomlab.github.io/flu-seqneut-2026/H1_prot_struct_viz.html#view=antigenic-regions for these regions projected on the HA structure and their constituent sites defined).

### Human sera and plasma

Details on the sera, including the collection date and age of the individual from whom it was obtained, are provided in **Table 1** and **Supplementary File 2**.

The VIDRL adult and elderly sera were collected by two sites with funding provided by Seqirus Limited for another study titled “A Sero-Epidemiological Study for the Collection of Pre- and Post-Vaccination Blood Samples from Volunteers Receiving a Southern Hemisphere Formulation of Inactivated Influenza Vaccine Recommended for the Season”; the de-identified sera reagents were then shared for the current study. Human research ethics approvals for this study were from the Adelaide University Women’s and Children’s Health Network Human Research Ethics Committee (application number 2020/HRE01220) and the University of Sunshine Coast Clinical Trials Bellberry Human Research Ethics Committee (application number 2025-12-2108).

The VIDRL child sera were collected from a single site with funding provided by AstraZeneca for another study titled “SNIF-FLES: Southern Hemisphere Nasal Influenza Flu Vaccine Experience Study”; the de-identified sera reagents were then shared for the current study. Human research ethics approval for this study was from Murdoch Children’s Research Institute, Melbourne (application number HREC/120054/RCHM-2026).

The SCH sera were obtained from children not known to be immunocompromised receiving medical care at Seattle Children’s Hospital, mostly for reasons unrelated to respiratory viral infection. The sera were obtained with a signed waiver of consent and the approval of the Seattle Children’s Hospital Institutional Review Board.

The CTS plasma samples were collected from volunteer adult blood donors across regions of the United States through a biobank maintained at Creative Testing Solutions by a collaboration between Vitalant Research Institute and the American Red Cross.

The deidentified remnant sera from UWMC were obtained from adults testing positive for HBsAb (to identify individuals with intact immunity) with approval from the University of Washington Institutional Review Board with a consent waiver.

All plasma and sera samples were processed identically before use in sequencing-based neutralization assays. Plasma samples were spun at 2000g for 10 minutes and cleared supernatant was transferred to new tubes. All sera and plasma were then treated with receptor-destroying enzyme II and heat-inactivated following a previously described protocol (Lee et al. 2019) to remove sialylated compounds that can neutralize virus by binding to the HA receptor-binding pocket, and to inactivate complement proteins. Briefly, lyophilized receptor-destroying enzyme II (Seiken) was resuspended in sterile phosphate-buffered saline (PBS) per manufacturer instructions and then passed through a 0.22 um filter before being aliquoted and frozen at −20C. These aliquots were thawed immediately prior to use, never undergoing more than a single freeze-thaw cycle. 25 uL of clarified plasma or sera was then incubated with 75 uL of receptor-destroying enzyme in 96-well PCR plates (Thermo Scientific) at 37C for 2.5 hours and then 55C for 30 minutes. All sera and plasma were used immediately following receptor-destroying enzyme II-treatment or stored at 4C until use. The titers reported here account for this initial four-fold dilution of the sera or plasma during this treatment.

### Choice of HAs for the sequencing-based neutralization assays

Our process for choosing HAs for the sequencing-based neutralization assay library aimed to select HAs that would be representative of circulating diversity in H1N1 and H3N2 HAs during the 2027 Southern Hemisphere influenza season. Due to the experimental time requirements for generating this library and running the neutralizations assays prior to the vaccine-selection meeting in September 2026, we made these selections largely based on data available in late May 2026. We first identified all HA protein haplotypes present in sequenced human influenza using the latest Nextstrain pdmH1N1 and H3N2 6-month builds available in May 2026 (Hadfield et al. 2018). We then created a ranked list of all haplotypes for each subtype based on a weighted combination of the following features: the total number of HA1 mutations, HA mutations in previously defined antigenically important sites or sites in/near the receptor binding site (Koel et al. 2013; Wolf et al. 2006; Caton et al. 1982), HA mutations arising at a rate greater than expected given the underlying mutation rate (Bloom and Neher 2023; Turner et al. 2026), the relative growth advantage of the haplotype compared with other concurrently circulating strains as defined by Nextstrain’s multinomial logistic regression forecast model (Abousamra et al. 2024), and the degree of geographic spread of the haplotype. The code for generating these lists of HA haplotypes is available at https://github.com/nextstrain/seasonal-flu/blob/fc2dd65/scripts/summarize_haplotypes_for_library_design.py We used these ranked lists along with expert selection and curation to choose an initial set of 77 H3N2 haplotypes and 63 H1N1 haplotypes. Documentation and logic describing the design of the barcoded HA constructs is at https://github.com/jbloomlab/flu-seqneut-2026/tree/main/non-pipeline_analyses/library_design/full_design.

Additionally, this library was designed to cover the recent component strains for both the H1N1 and H3N2 seasonal vaccine strains. For H3N2, we included cell- and egg-based vaccine strains from the 2024 Southern Hemisphere vaccine to the 2026-2027 Northern Hemisphere vaccine. For H1N1, we attempted to include cell- and egg-based vaccine strains from the 2020-2021 Northern Hemisphere vaccine to the 2026-2027 Northern Hemisphere vaccine. However, not all H1N1 egg-based vaccine strains within this time frame were included in the library design as several do not grow well in the mammalian cell culture system used for our experiments; specifically, the following H1N1 vaccine strains were not included: A/Guangdong-Maonan/SWL1536/2019 (egg), A/Victoria/2570/2019 (egg), A/Sydney/5/2021 (egg and cell), and A/Victoria/4897/2022 (egg).

Overall, this design process resulted in a set of 154 HAs (most of which are associated with multiple distinct barcodes) that were included in the original designed library (see https://github.com/jbloomlab/flu-seqneut-2026/blob/main/data/viral_libraries/flu-seqneut-2026-barcode-to-strain-designed.csv). The set of 148 HAs that were actually included in viruses that could be generated at sufficient titers for use in the sequencing-based neutralization assays is at https://github.com/jbloomlab/flu-seqneut-2026/blob/main/data/viral_libraries/flu-seqneut-2026-barcode-to-strain-actual.csv and in **Supplementary File 1**.

### Production and pooling of viruses for sequencing-based neutralization assays

The sequencing-based neutralization assays measure titers via barcode sequencing, requiring HA genes to be linked to nucleotide barcodes. As in prior work (Bacsik et al. 2023; Welsh et al. 2024; Kikawa et al. 2026b; Loes et al. 2024), we linked HA genes to 16-nucleotide barcodes without disrupting genome packaging by duplicating the region at the end of the coding sequence containing the packaging signals. Randomly generated 16-nucleotide barcodes were cross-checked to avoid barcode collision (with other barcoded HA constructs from prior libraries) and barcodes starting with ‘GG’ (which we have found sequence poorly). The barcoded HA genes were then cloned into a plasmid backbone by Twist Biosciences or Genscript. For both H3N2 and H1N1 HA plasmids we used either a derivative of the pHH21 unidirectional reverse genetics plasmid (Neumann et al. 1999) or the pHW2000 bidirectional reverse genetics plasmid (Hoffmann et al. 2000). See https://github.com/jbloomlab/flu-seqneut-2026/blob/main/non-pipeline_analyses/data/example_vRNA_eGFP.gb for a Gen-Bank file with the full annotated HA vRNA containing GFP in place of HA (used as the template for cloning), and https://github.com/jbloomlab/flu-seqneut-2026/blob/main/non-pipeline_analyses/data/example_vRNA_barcoded_H1_HA.gb or https://github.com/jbloomlab/flu-seqneut-2026/blob/main/non-pipeline_analyses/data/example_vRNA_barcoded_H3_HA.gb for example barcode HA vRNAs with a H1 or H3 HA ectodomain. While the bulk of the library was newly created, some constructs were generated in previous studies (Kikawa et al. 2025; Kikawa et al. 2026a; Loes et al. 2026; Kikawa et al. 2026b).

The barcoded HA plasmids were then used to individually generate barcoded virions using reverse genetics, which were then pooled together for use in sequencing-based neutralization assays. All HA genes were barcoded in duplicate or triplicate, and these replicate barcoded plasmids were pooled together prior to use. As previously described, each barcoded HA construct was transfected with all other non-HA genes from the lab-adapted A/WSN/1933 strain (see **Biosafety** subsection above) onto a co-culture of 293T and MDCK-SIAT1-TMPRSS2 and rescued 72 hours post transfection. These rescued viruses were then individually passaged on MDCK-SIAT1-TMPRSS2 cells.

The virus library pools used in sequencing-based neutralization assays should contain each barcoded virus at roughly equal proportions. As described previously, we first made an equal-volume pool of each individually generated and passaged virus and then titrated across MDCK-SIAT1 cells to determine each virus’s relative transcriptional titer (see report here: https://jbloomlab.github.io/flu-seqneut-2026/20260715_equal_vol_pool_analyze_pool.html). Using these estimated transcriptional titers, we repooled and re-titrated the pool to confirm the pool was more equally balanced and to determine the virus dilutions where viral transcription increased proportionally with the amount of virus infected on cells (see report here: https://jbloomlab.github.io/flu-seqneut-2026/20260810_repool_wash_spikein_lysis_measure_repool.html). Based on this analysis, the viral pool was used at a 1:64 dilution, since this was the highest virus pool concentration that was clearly in the linear range of viral barcode transcription as a function of infectious viral dose, as determined by dilution series analysis at https://jbloomlab.github.io/flu-seqneut-2026/20260810_repool_wash_spikein_lysis_measure_repool.html (the sequencing based readout requires transcription to scale linearly with viral infectivity).

The presence of virions containing different barcoded HAs in their genome than expressed on particle surface as a result of cross-contamination would be a major confounder for titers measured in the sequencing-based neutralization assay. To test for this type of cross-contamination, we individually infected each passaged virus on 50,000 MDCK-SIAT1 cells in a single well of a 96-well plate. At 16 hours post infection, we extracted viral RNA and sequenced per normal protocol for sequencing-based neutralization assays (see below). The results of that report are here (https://jbloomlab.github.io/flu-seqneut-2026/20260730_single_well_analyze_single_well_infections.html). This analysis confirmed none of the viral strains used in our experiments had appreciable contamination.

### Sequencing-based neutralization assays

The neutralization assays were performed as described previously (Kikawa et al. 2025; Kikawa et al. 2026a). The detailed protocol for these assays is available at https://doi.org/10.17504/protocols.io.kqdg3xdmpg25/v2; all details including the cell line, cell numbers, timepoints, etc are described in that protocol. As described in that protocol, infectivity is quantified by sequencing the viral barcodes in infected cells, and normalizing the barcode counts to an RNA standard consisting of known barcodes that is added at the cell lysis stage. Normalization to this RNA standard allows conversion of barcode sequencing counts to fraction viral infectivity retained at each serum concentration (Loes et al. 2024).

The initial serum dilution was 1:40 (accounting for the initial 1:4 dilution from the receptor-destroying enzyme treatment described above), making that the lower limit of detection for the assays. Matching plate formatting from prior iterations of this experiment (Kikawa et al. 2025; Kikawa et al. 2026a), sera were then diluted 2.3-fold across each 8-well column of a 96-well plate, making the upper limit of detection 1:13,620. The twelfth column of every 96-well plate was used for no-serum control wells.

### Analysis of sequencing-based neutralization assay data

We analyzed the sequencing-based neutralization assay data using *seqneut-pipeline* (https://github.com/jbloomlab/seqneut-pipeline), version 9.2.2. This pipeline uses *neutcurve* (https://github.com/jbloomlab/neutcurve) to fit neutralization curves to the fraction infectivity calculated from the barcode sequencing. It also applies quality control filters (configured at https://github.com/jbloomlab/flu-seqneut-2026/blob/main/config.yml) to remove low-quality neutralization curves. This filtering is why the final titer data set has 52,268 titers for the 148 viral strains measured against 355 sera rather than 148 × 355 = 52,540 titers: 272 curves were dropped during quality control. See https://github.com/jbloomlab/flu-seqneut-2026 for all computer code implementing the analysis, and https://jbloomlab.github.io/flu-seqneut-2026 that shows all measured curves for each plate and sera along with the full quality-control report.

All HA-expressing strains were barcoded in duplicate or triplicate (as described above), but in the analysis for this study we collapsed all barcode counts for the same HA by summing the counts—this is a different analysis approach than our prior studies (Kikawa et al. 2025; Kikawa et al. 2026a; Kikawa et al. 2026b; Loes et al. 2024), where we analyzed each barcode separately and then took the median of the measured titer for each barcode for the same strain. The potential benefit of analyzing each barcode for a strain separately is that the per-barcode replicate measurements provide multiple technical replicates for each HA neutralization curve. However, a primary source of experimental noise in the assay is that only a finite number of infectious particles for each variant are added to each well, and this number is two- to three-fold lower if variants are analyzed at the barcode rather than the strain level (since there are two to three barcodes per strain). Empirically we assessed that any benefits from having multiple barcode replicates for each strain were outweighed by the increased noise due to fewer infectious particles per variant per well when the data are analyzed on a per-barcode rather than per-strain level. Therefore, we added a collapse_strain_barcodes flag to the analysis pipeline and analyzed each strain as the sum of its barcode counts. See *seqneut-pipeline* documentation on the collapse_strain_barcodes flag for more details (https://github.com/jbloomlab/seqneut-pipeline/#collapse_strain_barcodes); note the flag’s value can be reversed in the main analysis pipeline to re-analyze the data on the barcode level, although we assess for this study that collapsing barcodes gives better-quality data.

For the figures in this paper, we show titers to all recent human strains and cell-based vaccine strains that were used as far back as the 2024 Southern Hemisphere influenza season. The figures in this paper do not show the titers to the egg-based vaccine strains or older cell-based vaccine strains. The reason is that we tend to measure higher neutralization titers of egg-based vaccine strains (Kikawa et al. 2026b) and older cell-based vaccine strains, so including them in the figures can “blow out” the scale and obscure the more relevant differences among recent strains. However, the titers to older vaccine strains and egg-based vaccine strains are plotted at https://jbloomlab.github.io/flu-seqneut-2026/summary.html#titers-to-older-and-egg-based-vaccine-strains and are provided in numerical form in the supplementary files and CSVs on the GitHub repository.

### Phylogenetic trees and subclade nomenclature

The phylogenetic trees available at https://nextstrain.org/community/jbloomlab/flu-seqneut-2026@main/H3N2 and https://nextstrain.org/community/jbloomlab/flu-seqneut-2026@main/H1N1 and shown along the x-axis of some figures in this paper are built on the HA protein sequences of the strains included in the library, and have branch lengths that represent the number of amino-acid mutations. See https://github.com/jbloomlab/nextstrain-prot-titers-tree (which is included as a git submodule for the main project GitHub repository) for the code used to build these trees.

On the trees and in the figures, strains are labeled by their sub-clade from the nomenclature described in (Neher et al. 2026) plus any additional HA amino-acid mutations. Any mutations falling in the HA2 region are annotated as such, whereas mutations in the HA1 region have no additional annotations. For instance, K:V223I means the strain has the subclade K HA amino-acid sequence plus the V223I mutation in HA1, whereas K:Q80K,HA2_V77I means the strain has subclade K amino-acid sequence with Q80K in HA1 and V77I in HA2.

## Supplementary Files

**Supplementary File 1. Details about viral HAs used in this study**. This CSV is also available at https://github.com/jbloomlab/flu-seqneut-2026/blob/main/results/final_titer_data/human_viruses.csv

**Supplementary File 2. Details about sera used in this study**. This CSV is also available at https://github.com/jb2026/blob/main/results/final_titer_data/human_viruses.csv

**Supplementary File 3. Neutralization titers measured in this study. This CSV is also available at https://github.com/jbloomlab/flu-seqneut-2026/blob/main/results/final_titer_data/human_titers.csv**

